# Optimal MR-ARFI encoding parameters for a range of contemporary magnetic field gradient systems

**DOI:** 10.64898/2026.10.13.758785

**Authors:** Shota Hodono, David G. Norris

## Abstract

**Purpose:** To optimize motion encoding gradients in MR-Acoustic Radiation Force Imaging (ARFI) experiments under a range of experimental conditions, and to provide a general simulation tool.

**Methods:** In MR-ARFI, Transcranial focused Ultrasound Stimulation (TUS) induced tissue displacement is encoded by motion encoding gradients as a phase change. We extended previously proposed equations for phase SNR to account for finite gradient switching times. We simulated the phase SNR while varying the strength and slew rate of motion encoding gradients for monopolar and bipolar encoding schemes in order to identify optimal configurations for various gradient hardware specifications. Phantom MR-ARFI experiments were performed to validate the simulations.

**Results:** Optimized gradient durations are derived for a range of gradient strengths and slew rates. Gradient slew-rate is shown to influence the optimal parameters. Phantom MR-ARFI results showed good agreement with the simulations.

**Conclusion:** The availability of modern gradient hardware is shown to dramatically improve the sensitivity of the ARFI experiment. Our openly available calculation tool allows ready calculation of optimal timings for any experimental conditions.

## Introduction

Magnetic Resonance-Acoustic Radiation Force Imaging (MR-ARFI)^1–3^ is currently the only non-invasive technique capable of measuring subtle macroscopic displacements caused by low intensity transcranial focused ultrasound stimulation (TUS). Following a first successful implementation in humans^4^ it can safely be assumed that there will be a boom in the use of ARFI for planning and monitoring TUS-MRI applications.^5^

MR-ARFI measures a phase change that is proportional to the mean displacement during sonication. This is achieved by using gradient-encoding schemes similar to those employed in Diffusion Weighted Imaging (DWI). In a unipolar gradient scheme TUS will typically be applied during one of the two gradient lobes, and in a bipolar scheme during two out of four lobes.^6^ It was previously shown that a bipolar scheme is more sensitive than a unipolar scheme, primarily because of the reduced signal attenuation due to diffusion.^6^ It is also clear that the sensitivity of the ARFI experiment is strongly dependent on the maximum gradient strength available.

Driven by the benefits for DWI, there is a continuing effort within the MRI community to increase the strength of the gradient field available. This has led to ever-increasing gradient strengths for whole-body systems, whereby the slew rate is largely constrained by physiological considerations to about 200-250 T/m/s, and to the development of a range of dedicated head-only gradient sets where the physiological constraints are lower and the smaller inner diameter facilitates stronger maximum gradients.

In the interests of optimising experimental conditions and harmonisation, we here extend the formulation given in reference^6^ to include finite slew rates, and to present optimal ARFI acquisition parameters for a range of current and future gradient systems. The validity of the formulation is experimentally verified using a 3T system.

## Methods

MR-ARFI employes a spin-echo sequence with motion encoding gradients (MEG)^2^. As Figure 1 shows, by synchronizing TUS with MEG, the TUS-induced tissue displacement is encoded as a phase change. The accrued displacement induced phase, *ϕ*, is expressed by

**Figure 1.**
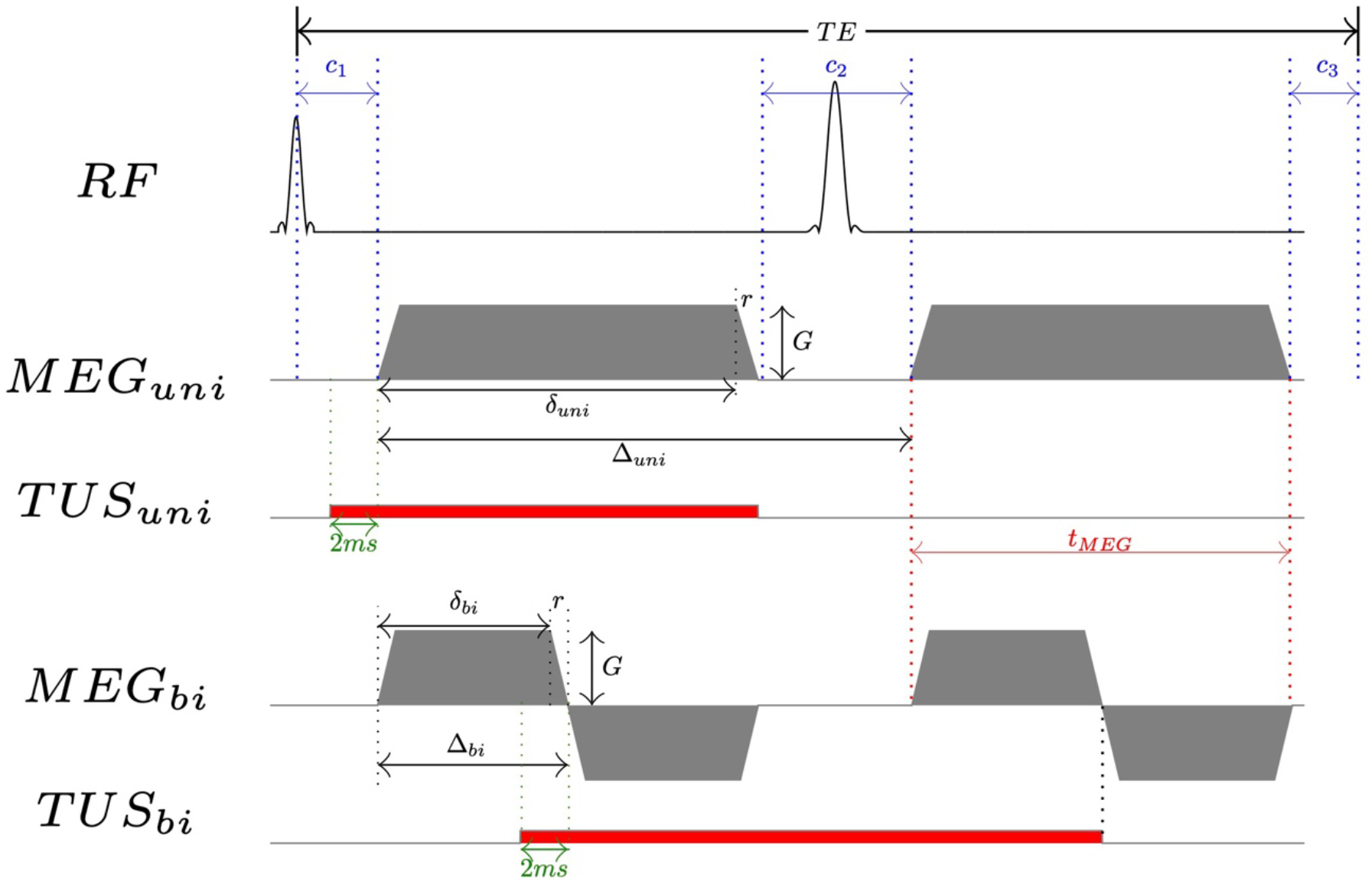
MR-ARFI sequence diagram for unipolar and bipolar encoding schemes. δ is defined as the sum of the flat-top and ramp-up times, and Δ is the time between the initial rise of the first and second motion encoding gradient pulses. The ramp-up and ramp-down time are defined as r. The parameters c_1_, c_2_, and c_3_ represent the time intervals from the center of the excitation pulse to the start of the first gradient pulse, between the end of the first gradient pulse and the start of the second, and from the end of the second gradient pulse to the echo time (TE), respectively. The red squares represent TUS duration. To account for the tissue-displacement rise time^11^, TUS was turned on 2 ms^4^ before the encoding gradients.

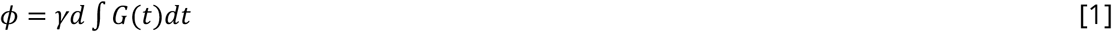

where the γ, d, and G are the gyromagnetic ratio, tissue displacement, and gradient amplitude. In general, TUS ‘on’ and ‘off’ volumes are acquired then subtracted to experimentally compute TUS-induced phase change, Δϕ. With that, displacement can be computed by

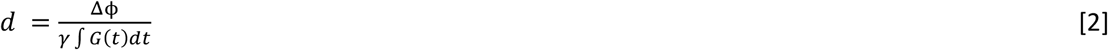

The equations given in Chen et al.^6^ for the SNR of the MR-ARFI experiment using unipolar and bipolar displacement encoding are respectively

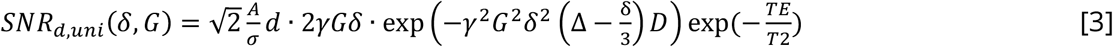

and

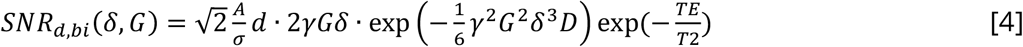

where A, σ, and δ are the signal intensity in the absence of noise, the standard deviation of the Rician phase noise,^7^ and motion encoding gradient duration for one lobe in unipolar scheme.

By reference to figure 1 we hereby extend these equations, using

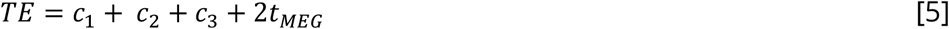

where c_1_, c_2_, and c_3_ are preparation time, and *t*_*MEG*_ is the gradient duration of single lobe and two lobes in unipolar and bipolar schemes respectively;

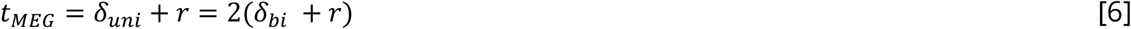

By including the effect of finite switching times on the displacement induced phase change and the b-value, equations 3 and 4 become

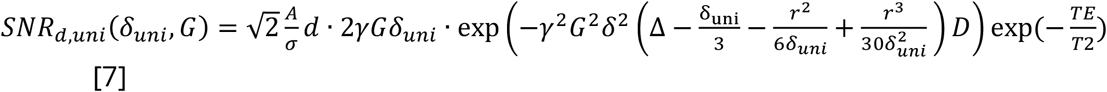

for unipolar gradients^8,9^. Where

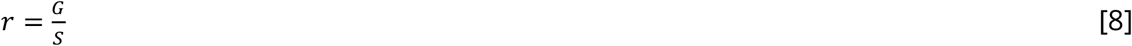

where S is the slew rate.

And

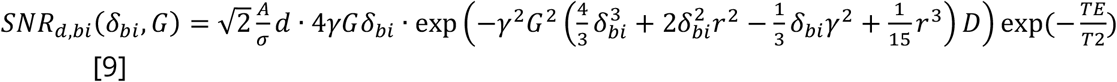

for bipolar. Note that, in the bipolar scheme, second and forth gradient lobes were started without delay.^10^

### Simulations

In this work, we simulated equation [7] and [9] to investigate the relationship between the phase SNR and motion encoding gradient strengths and slew rates for unipolar and bipolar schemes. The gradient duration (*t*_*MEG*_), strength, and slew-values were varied from 4 to 30 ms, 20 to 300 mT/m, and 100, 200 and 600 T/m/s respectively, with all other parameters fixed (ADC = 0.9 × 10^-9^ m^2^/s^12^ and T2 = 80 ms^13^ for gray matter at 3T). In addition, we specifically considered some commonly available gradient sets across vendors. The code used for the simulations is available at GitHub (https://github.com/s-hodono/optARFI). In the simulations the following parameters were used. Note that, as seen in equation [7] and [9], TE is varied according to *t*_*MEG*_ while fixating preparation time.

### Experiments

To validate the simulations, phantom MR-ARFI experiments were performed sweeping *t*_*MEG*_and G in unipolar and bipolar schemes using Siemens Prisma 3T MRI. A gelatine phantom made with evaporated milk^14^ in a 3D printed container (Figure 2) was scanned at 3T using the standard 20 channel head-neck coil (Magnetom Prisma, Siemens Healthineers, Germany). 300 kHz annular array transducers (IMASONIC, France) were used and driven using a 32 channel system^15^ (Image Guided Therapy, Pessac, France). The complete assembly was submerged in a de-gassed water-filled container. TUS focal point was set to 65 mm and output power in water was 1.0 MPa in all experiments. The TUS duration was varied according to *t*_*MEG*_.

**Figure 2.**
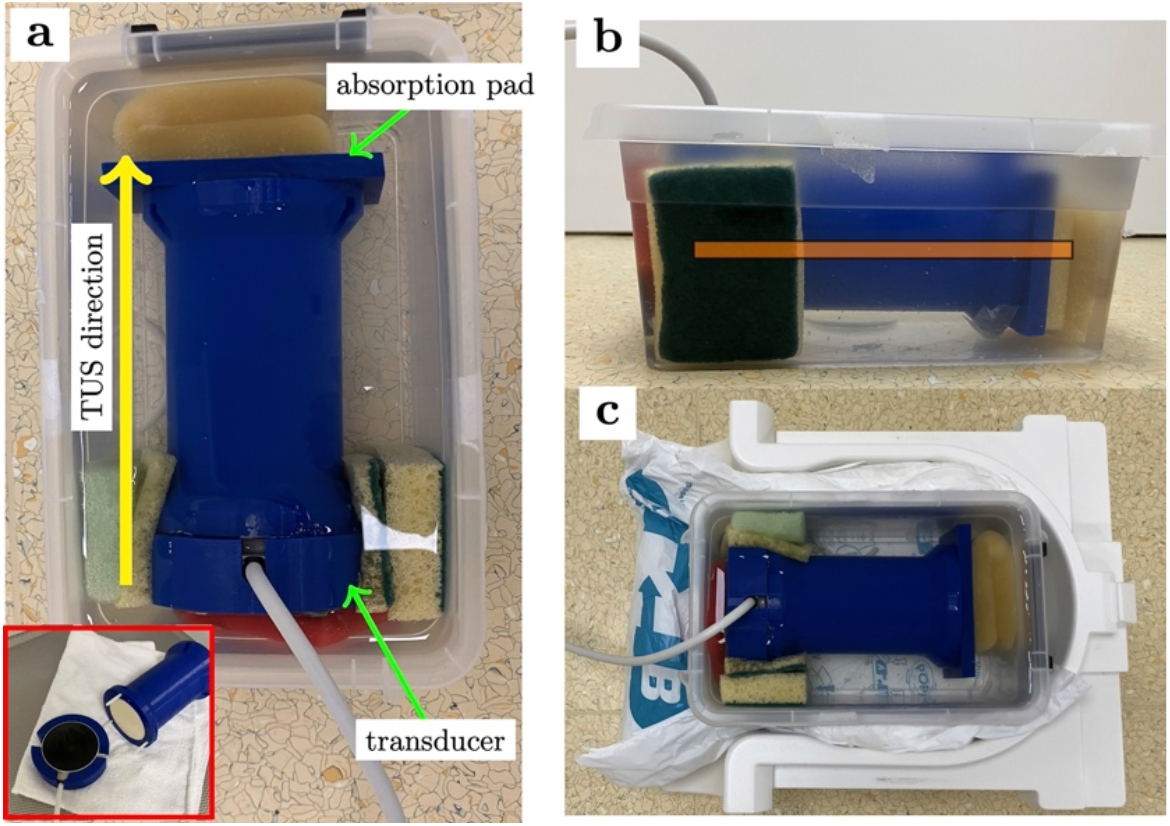
MR-ARFI experimental setup. (a) A tube filled with gelatine phantom was submerged in water with a transducer. The small red window shows phantom surface and transducer. (b) imaging slice is visualized as an orange rectangular box. (c) Phantom placed in 20Ch head-neck coil. Anterior part of the coil was placed after positioning the phantom.

The MR-ARFI pulse sequences were prepared using *pulseq*^16^ based on a diffusion weighted -HASTE sequence.^17^ The sequence parameters are as follows: number of dummy cycle = 4, slice TR = 3 s, matrix size = 80 × 72, FOV = 240 × 216 mm^2^, slice thickness = 3 mm, number of volumes = 20 (TUS ‘on’ and ‘off’ volumes were alternated), intra-echo TE = 7 ms, preparation time (c_1_ + c_2_ + c_3_) = 46 ms. For unipolar and bipolar experiments, an identical readout was used but different gradient schemes (Figure 1). The gradient duration (*t*_*MEG*_) and G were varied from 4 to 30 ms and 20 to 75 mT/m. In the bipolar scheme, the motion encoding gradients were played-out in motion-compensated manner. In order to avoid stimulation limits errors related to the gradient switching, the slew rate was varied across experiments. The slew rate used for the experiment is seen in Supplementary Figure 1.

To compare the experimental results to the simulations, MR-ARFI phase changes were averaged in an ROI drawn on the bipolar MR-ARFI image acquired with *t*_*MEG*_ = 12 ms and G = 75 mT/m (see Supplementally Figure 2). The equation [7] and [9] account SNR and phase encoding efficiency. Therefore, temporal SNR (tSNR) and MR-ARFI phase changes (Δϕ) were computed in the ROI and multiplied to compare to the simulations.

## Results

### Simulations

Figure 3 shows simulation results for unipolar and bipolar schemes with different slew rates, 100T/m/s, 200T/m/s, and 600T/m/s respectively. Clearly, the bipolar scheme shows higher sensitivity as previously observed^6^ and higher slew rates contribute to higher sensitivity. Table 1 summarizes optimal parameters for each standard gradient configuration and relative SNR gain compared to a basic clinical scanner (Gmax = 45mT/m, *t*_*MEG*_ = 10ms with S = 200 T/m/s). As expected, using the maximum available gradient amplitude will maximize the phase SNR, the simulations show how the optimal gradient duration varies with gradient strength and slew rate.

**Table 1.** Vendor specific gradient specs and the optimal ARFI parameters. In all scenarios, bipolar scheme showed higher sensitivity. The gradient duration, *t*_*MEG*_, is a total duration of two gradient lobes (see Figure 1). *As impulse gradient system is available at 7T, T2 = 45ms was used for the simulation.

| vendor | system | type | Max<br>gradient<br>amplitude<br>[mT/m] | Max<br>slew<br>rate<br>[T/m/s] | Optimal parameters |  |  | Relative<br>phase<br>SNR<br>gain |
| --- | --- | --- | --- | --- | --- | --- | --- | --- |
| | | | | | Gradient<br>duration<br>($t_{MEG}$) [ms] | $\delta_{bi}$ [ms] | Gradient<br>strength<br>[mT/m] | |
| Siemens | Skyra | whole<br>body | 45 | 200 | 21.4 | 10.5 | 45 | 1.0 |
|  | Prisma | whole<br>body | 80 | 200 | 16.2 | 7.7 | 80 | 1.4 |
|  | Cima. X | whole body | 200 | 200 | 10.6 | 4.3 | 200 | 2.1 |
|  | Connectome 2.0 | Head only | 500 | 600 | 6.4 | 2.4 | 500 | 3.1 |
|  | Impulse gradient* | Head only | 200 | 900 | 8.9 | 4.2 | 200 | 1.7 |
| Philips | Ingenia | whole body | 45 | 200 | 21.4 | 10.5 | 45 | 1.0 |
|  | Ingenia Elition X | whole body | 65 | 220 | 17.9 | 8.7 | 65 | 1.3 |
|  | Titanion |  | 150 | 250 | 11.7 | 5.3 | 150 | 1.9 |
| GE | Architect | whole body | 44 | 200 | 21.6 | 10.6 | 44 | 0.99 |
|  | Premier | whole body | 80 | 200 | 16.2 | 7.7 | 80 | 1.4 |
|  | MAGNUS | Head only | 300 | 750 | 7.7 | 3.5 | 300 | 2.7 |

**Figure 3.**
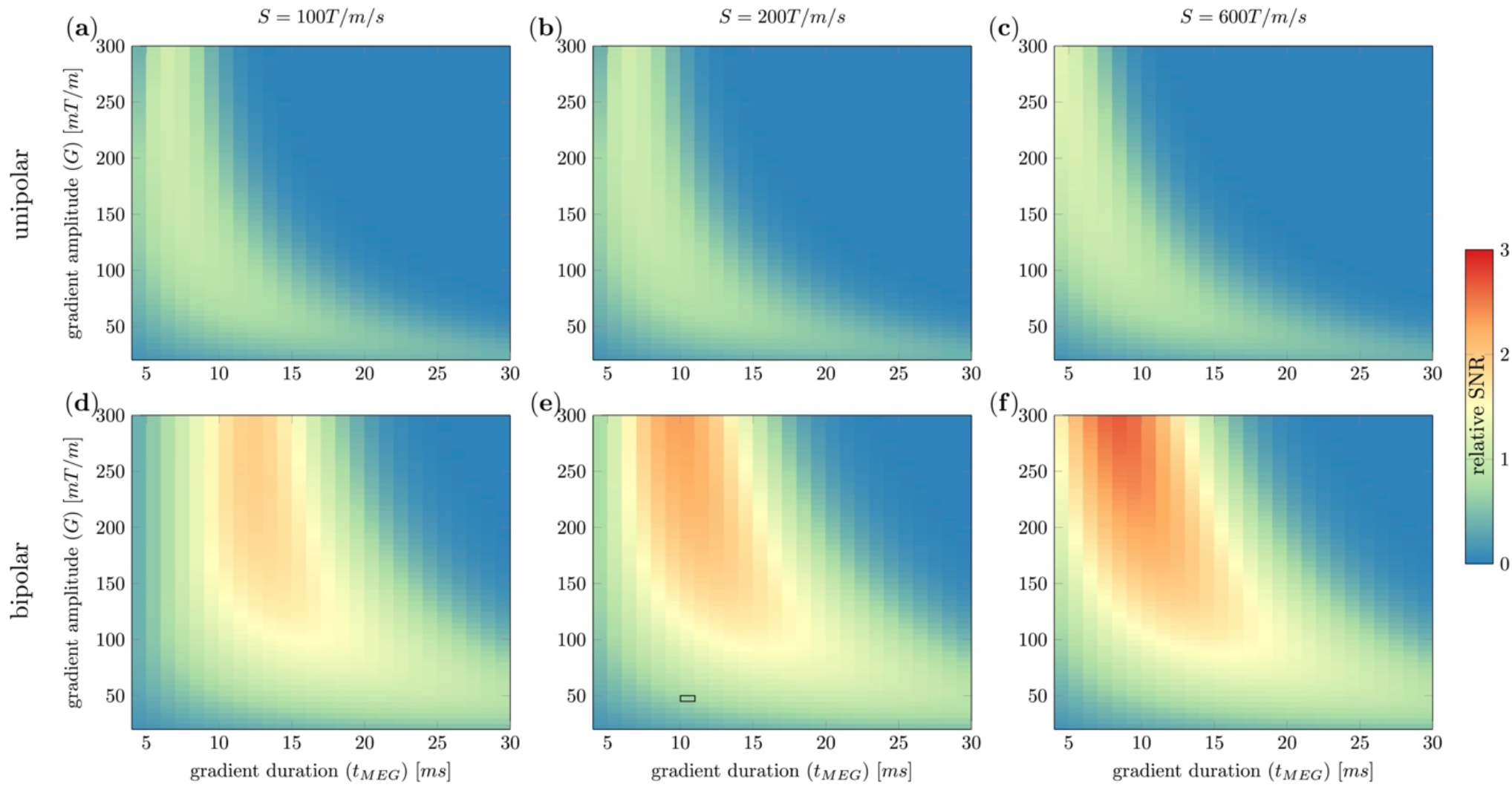
Simulation results of MR-ARFI sensitivity, (**a, b, c**) unipolar with slew rate of 100, 200, and 600 T/m/s, (**d, e, f**) bipolar with slew rate of 100, 200, and 600 T/m/s. All simulations were normalized by the value acquired with G = 45 mT/m, *t*_*MEG*_ = 10ms, and S = 200 T/m/s, highlighted by the black rectangle in panel (**d**). All simulations are plotted in same color limits.

### Experiments

Figure 4 shows simulated phase SNR using the equation [7] and [9] and experimental results. The experimental results showed similar trend to the simulations in both unipolar and bipolar schemes. Individual ARFI phase change images and tSNR images were seen in supplementary figure 3.

**Figure 4.**
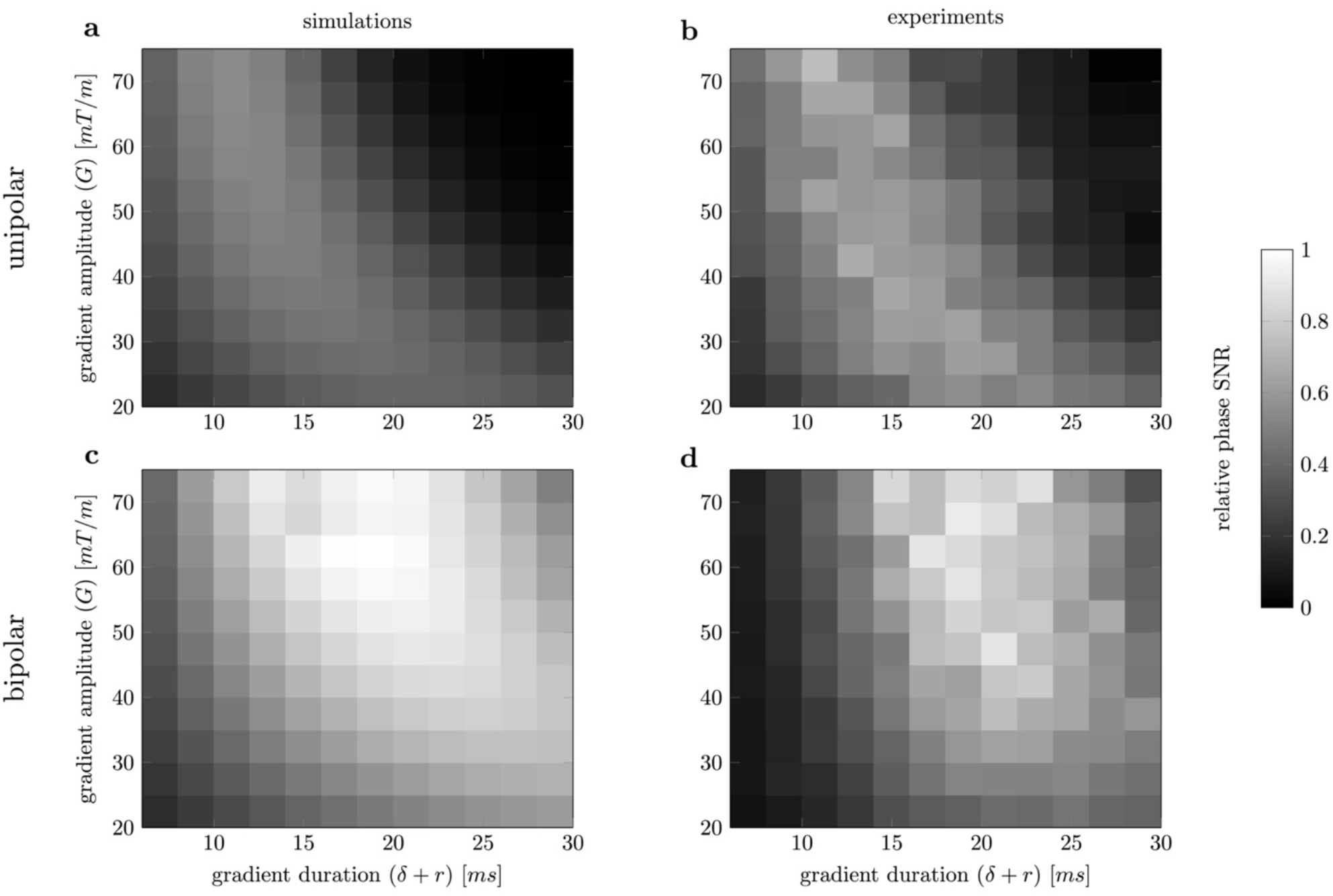
Phase SNR: simulations vs experiments. (**a, c**) simulated phase SNR in unipolar and bipolar schemes. (**b, d**) measured phase SNR in unipolar and bipolar schemes. In this simulations, T2 = 200 ms, D = 1.5×10^-9^ m^2^/s, and d = 2 μm were used. Both simulations and experimental results were normalized by the highest SNR respectively.

## Discussion/Conclusion

Leveraging high-performance gradient system, ARFI contrast can be improved significantly. As previously shown^6^ bipolar schemes will always give higher phase SNR as the phase-change per unit displacement is associated with a lower b-value.

As higher gradient amplitude is used, the optimal gradient duration becomes shorter, and consequently, the required TUS duration also decreases. A lower TUS duty cycle reduces energy deposition at the TUS focal point, which is desirable to minimize unwanted neuromodulatory effects during MR-ARFI experiment and to reduce tissue heating.

This work showed that MR-ARFI phase SNR can be optimized by selecting highest available gradient amplitude with optimal gradient duration using bipolar scheme. Leveraging the high-performance gradient systems phase SNR can be enhanced significantly.

## Acknowledgements

This work was supported by the European Union’s Horizon Europe research and innovation program under grant agreement No 101071008. The authors acknowledge the FUS initiative at Radboud University.

