## Supplementary Figures for "Optimal MR-ARFI encoding parameters for a range of contemporary magnetic field gradient systems"

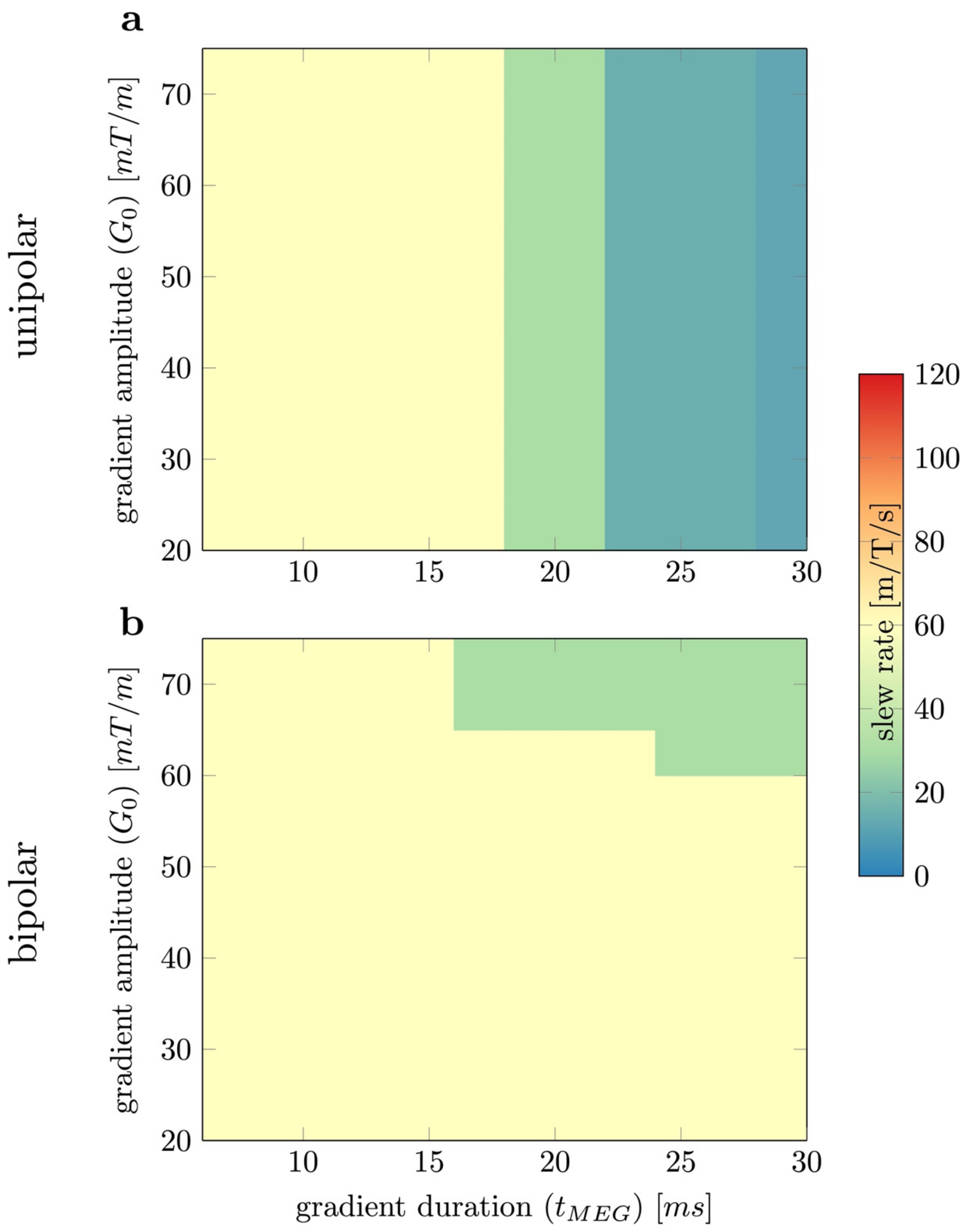


Supplementally Figure 1. Slew rates used in the experiments. In the experiment, very low slew rates were used to avoid stimulation errors.

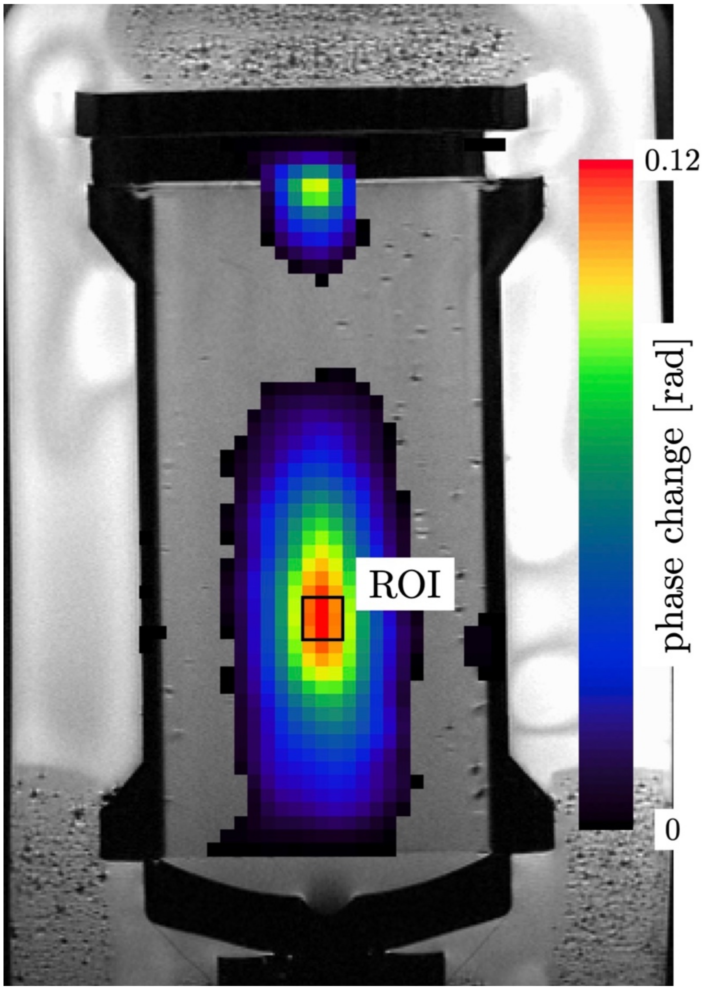


Supplementally Figure 2. ROI drawn on MR-ARFI image. The MR-ARFI phase difference image was acquired with $t_{MEG}$ = 12 ms and G = 75 mT/m using bipolar scheme. The MR-ARFI image was overlayed on high resolution T2 weighted structural image.

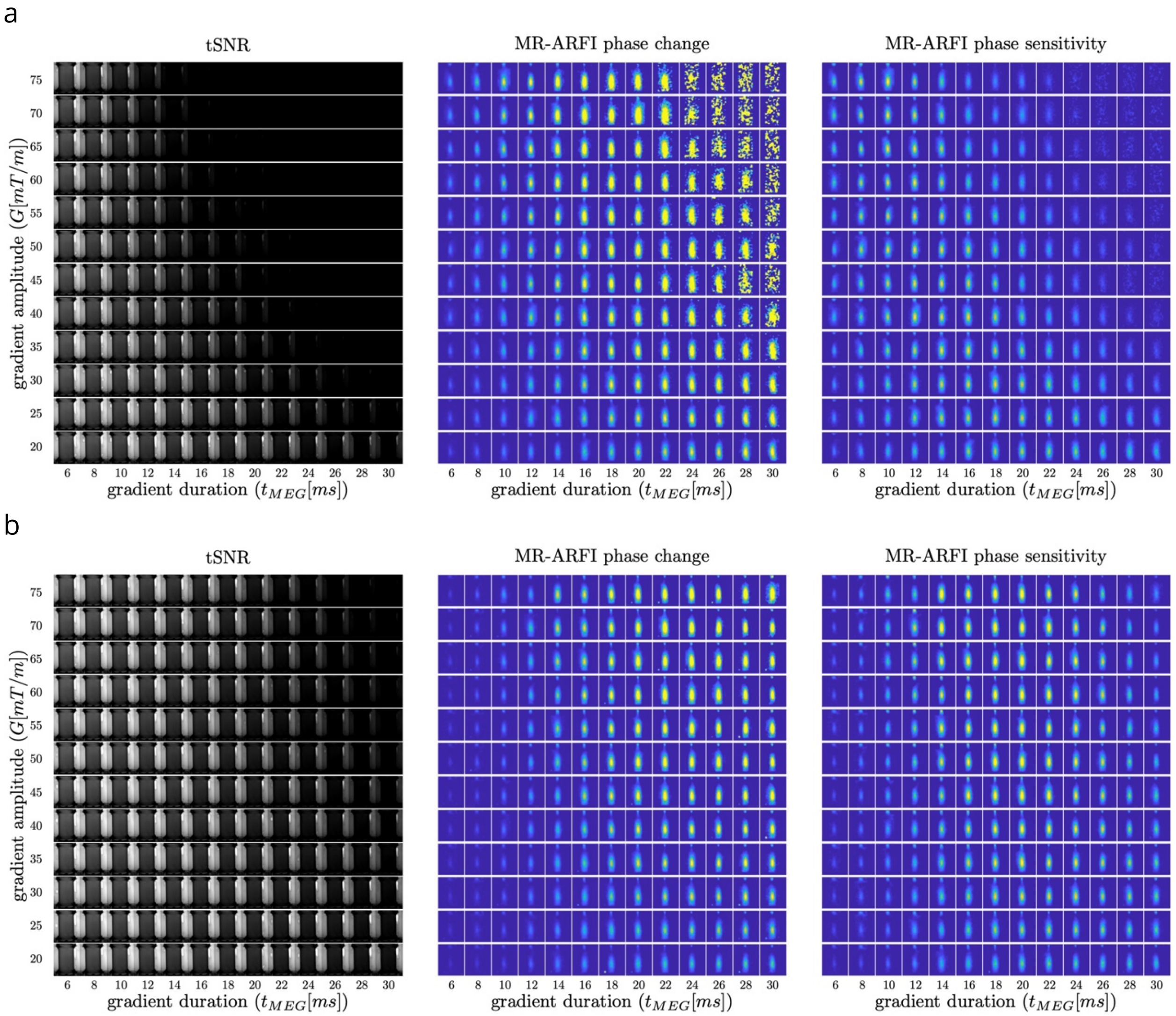


Supplementally Figure 3. tSNR, MR-ARFI phase change and sensitivity maps, a: unipolar, b: bipolar. MR-ARFI phase sensitivity images were acquired by multiplying tSNR and MR-ARFI phase change images.
